# A plasma metabolomics workflow for breast cancer detection using quantitative GC/MS and machine learning

**DOI:** 10.64898/2026.09.23.753710

**Authors:** Noriyuki Ojima, Tomonori Yokoyama, Shin Nishiumi, Shojiro Kikuchi, Kazue Yoneda, Misato Kimura, Kaisei Tanaka, Mai Morimoto, Ayako Miyazaki, Toshihiko Tomita, Mitsumasa Ohyanagi, Masayuki Nagahashi, Yasuo Miyoshi, Masataka Ikeda

## Abstract

Blood-based metabolomic profiling has been widely investigated for breast cancer (BC) detection; however, clinical implementation remains limited due to variability in sample handling, analytical reproducibility, and overfitting during statistical analysis. We established a plasma GC/MS metabolomics workflow for discriminating BC from healthy controls (HC) using conventional machine-learning algorithms. Plasma samples (n = 360; BC = 180, HC = 180) were collected prospectively under standardized preanalytical conditions before surgery and the initiation of systematic anticancer therapy and analyzed using a quantitative GC/MS platform with automated derivatization. Feature selection and model development were conducted using three machine-learning (ML) algorithms (Lasso logistic regression (LR), random forest classifier (RFC), and support vector machine (SVM)). A total of 45 metabolite candidate biomarkers were identified, and the optimal number of metabolite features for each algorithm was estimated by a recursive feature elimination (RFE)-based strategy. The best-performing models achieved area under the ROC curve values (AUC) of 0.910 (LR), 0.893 (RFC), and 0.843 (SVM). We selected prioritizing candidate biomarkers consistently expressed across the multi-algorithm pipeline. A bagging ensemble model improved stability (AUC = 0.911) and reduced false-positive predictions in the independent HC dataset. In addition, model stability with respect to false-positive predictions was assessed using an independent HC cohort (n = 15) that was collected at a separate institution. These results indicate that a plasma metabolomics workflow combined with conventional multi-algorithm ML, algorithm-specific feature selection, and independent assessment provides stable discrimination between BC and HC in a moderately sized cohort.

## Introduction

Metabolic changes occur downstream of genetic and proteomic regulation. As the phenotype closest to the biological state among the “-omics” layers, the metabolome provides insight into responses to internal and external perturbations, including growth, disease, genetic modification, and environmental factors [1]. Thus, metabolic profiling is a promising strategy for monitoring dynamic biological changes. Cancer is not only caused by congenital abnormalities, but also by acquired effects, in which various genetic abnormalities are caused by external factors, such as the living environment [2–6]. Therefore, cancer is an important target for studies using classification model-based metabolic profiling.

Breast cancer (BC) ranks fourth overall in mortality and first among women [7]. It generally progresses relatively slowly, and there is a marked difference in five-year survival rates between early- and advanced-stage disease. Early detection increases the likelihood of breast-conserving therapy, which contributes to improved survival and a better postoperative quality of life. Thus, BC is a malignancy for which early detection is particularly important. Consequently, BC has been extensively studied for biomarker discovery and classification model development by metabolic profiling [8, 9].

Most metabolomic studies have used nuclear magnetic resonance (NMR) [10, 11] and liquid or gas chromatography coupled with mass spectrometry (LC/MS or GC/MS) as analytical platforms. Because MS is more sensitive than NMR, it is commonly used for the analysis of low-abundance metabolites in liquid biopsy samples. LC/MS supports high-throughput analyses. When combined with high-resolution mass spectrometry, it enables targeted and non-targeted analyses [12–14]. GC offers a broader detectable metabolite range in a single assay compared with LC, and it is less affected by matrix effects and ion suppression from co-eluting compounds, which results in higher resolution [9, 15, 16].

Although numerous metabolomics studies on BC have been carried out with the aim of clinical application, there are few examples of metabolite biomarkers that have actually been implemented in clinical tests [8, 17, 18]. Such barriers to progress clinical tests include:

1. instability of metabolites depending on sample pretreatment and storage conditions [19];
2. issues with analytical reproducibility due to inter-institutional variability [17, 20];
3. risk of overfitting arising from the interaction among sample size, measurement noise, feature dimensionality, and model complexity; and
4. insufficient evaluation using samples independent of model development.

The present study has two complementary strengths to overcome these challenges.

First, with respect to sample quality, we conducted prospective sample collection suitable for metabolomics and established a GC/MS-based quantitative analytical method with less inter-institutional variability. Second, with respect to model assessment, we developed a classification model by comparing three conventional machine-learning algorithms using repeated cross-validation, feature elimination, and independently split test data, and performed an independent assessment of false-positive rate using healthy control samples collected at other facilities.

The primary contribution of this study is the construction and evaluation of a reproducibility-oriented plasma metabolomics workflow in a moderately sized cohort, together with an independent assessment of false-positive stability. This design enabled us to determine the extent to which conventional ML methods can extract a stable breast cancer-associated signal from quantitatively measured plasma metabolites.

We hypothesized that careful control of preanalytical and analytical variability, combined with restrained feature selection and independent assessment, can reduce overfitting and improve the practical reliability of plasma metabolomics-based breast cancer discrimination.

## Materials and methods

### Subjects and study design

This study was approved by the Ethics Committee of Hyogo Medical University (No. 3258) and was conducted in accordance with the Declaration of Helsinki. All participants provided written informed consent before participating in this study.

The design and workflow are presented in Fig 1. Patients were recruited from Hyogo Medical University Hospital from December 2019 to March 2025. Resected specimens were pathologically classified according to the 8^th^ edition of the Union for International Cancer Control TNM Classification of Malignant Tumors. Healthy volunteers for Workflow 1 and 2 were recruited from the Health and Medicine Clinic of Hyogo Medical University from September 2019 to March 2025. Healthy volunteers for Workflow 3 were recruited from Umeda Health and Medicine Clinic of Hyogo Medical University from July to August 2025.

**Fig 1.**
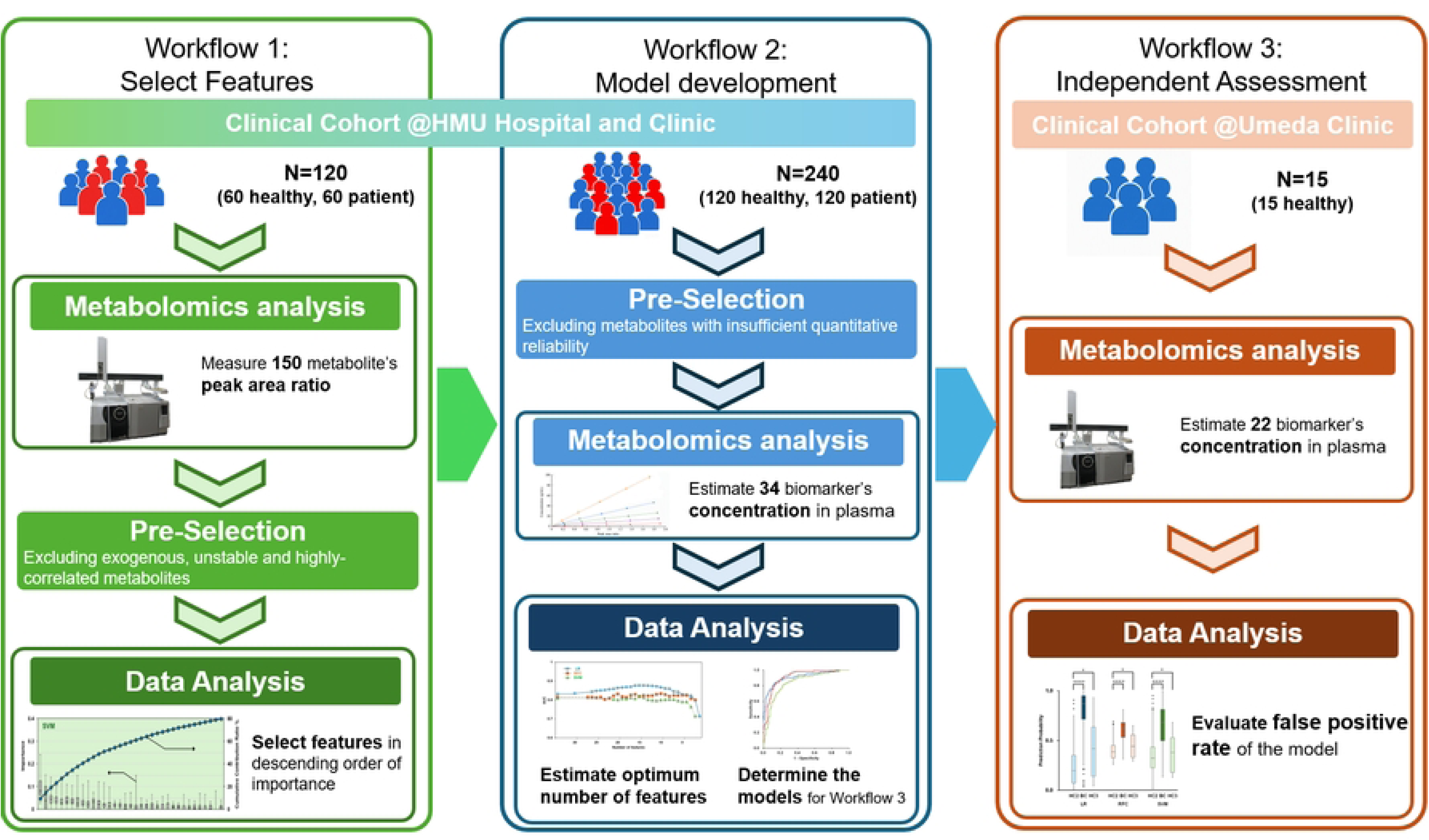
Schematic diagram of the study design. Workflow 1: Important metabolite features were selected from 150 metabolites using three types of machine-learning (ML) algorithms: LR, RFC, and SVM. Workflow 2: Estimate the optimum number of features for each ML algorithm and determine the model for Workflow 3 independent assessment. Workflow 3: 15 Healthy controls were collected independently of Workflow 1 and 2. The false-positive rate of the models was evaluated.

All patients and healthy volunteers who had a history of another advanced type of cancer within ten years, had previously been diagnosed with inborn errors of metabolism, thyroid abnormalities, hepatic or renal dysfunction, were pregnant or possibly pregnant, or had a history of primary liver cancer, were excluded. All patients were treatment-naive at the time of blood collection. Table 1 summarizes the characteristics of blood sample donors for Workflow 1, 2, and 3. A total of 180 BC and 180 HC blood samples were used for Workflows 1 and 2. The cancer patients were classified into 5-year age groups (30–34, 35–39, 40–44, …, 65–69, 70 or older). Healthy volunteers were selected by matching them as closely as possible with the cancer patients based on age. For Workflow 3, 15 HC samples were collected, and all individuals received breast mammography tests and ultrasonography.

**Table 1.** Characteristics of the blood sample donors.

|  | Workflow 1 |  | Workflow 2 |  | Workflow 3 |
| --- | --- | --- | --- | --- | --- |
|  | HC1 | BC1 | HC2 | BC2 | HC3 |
| **n** | 60 | 60 | 120 | 120 | 15 |
| **Age** |  |  |  |  |  |
| Mean | 56.1<sup>a</sup> | 57.7<sup>a</sup> | 55.9<sup>a</sup> | 57.2<sup>a</sup> | 53.3 |
| ±SD | 9.7 | 11.9 | 9.6 | 11.0 | 10.4 |
| **Gender** |  |  |  |  |  |
| Male | 0 | 0 | 0 | 0 | 0 |
| Female | 60 | 60 | 120 | 120 | 15 |
| **Stage** |  |  |  |  |  |
| 0/I/II/III/IV |  | 13/27/14/6/0 |  | 28/48/30/11/3 |  |

Abbreviations: BC, breast cancer, and HC, healthy controls.
<sup>a</sup>There is no significant difference between BC and HC by Wilcoxon's t-test.

### Blood collection

Subjects were only allowed to take water or Japanese tea for 8 hours before blood collection. Approximately 7 mL of unstimulated blood was collected in a blood collection tube (EDTA-2Na). After gentle mixing, the tubes were transferred within 1 min to a CubeCooler (Forte Grow Medical, Co., Ltd., Tochigi, Japan), which was stored at −20°C for at least 10 hours before collection and maintained at the appropriate temperature based on the manufacturer’s instructions to prevent degeneration of the hematologic metabolites [20, 21]. Within 3 hours after blood collection, the blood was centrifuged at 3,000 rpm for 15 min at 4°C, and 3 mL of plasma sample was obtained. The plasma samples were transferred to a clean tube and stored at −80°C.

### Chemicals and reagents

The human plasma used for quality control was purchased from Kohjin Bio Co. (Saitama, Japan). 2-Isopropylmalic acid and methoxyamine hydrochloride were obtained from Sigma-Aldrich Japan (Tokyo, Japan). N-Methyl-N-trimethylsilyltrifluoroacetamide (MS-TFA) was purchased from GL Sciences, Inc (Tokyo, Japan). Methanol and pyridine were purchased from FUJIFILM Wako Pure Chemical Co (Osaka, Japan). A standard alkane series mixture (C7 to C33) was obtained from Restek Co (PA, USA). Stable isotope-labeled 2-hydroxybutylic acid was purchased from CDN Isotopes (Quebec, CA). Other stable isotope labeled metabolites were purchased from Cambridge Isotope Laboratories, Inc (MA, USA).

### Plasma preparation and metabolomics analysis

To extract metabolites from plasma, 50 µL of each plasma sample was mixed with 270 µL of methanol containing internal standards. The mixture was shaken for 10 sec at room temperature and stored on ice for 10 min, followed by centrifugation at 20,000 g for 10 min at 4°C. Next, 200 µL of the obtained supernatant was transferred to a fresh tube and subjected to centrifugal evaporation and freeze-drying overnight. For oximation, 60 µL of 20 mg/mL methoxyamine hydrochloride dissolved in pyridine was added, the tube was sonicated for 20 min, and shaken at 1,200 rpm for 90 min at 30°C. The mixture was centrifuged at 20,000 g for 10 min at 20°C, and 40 µL of the supernatant was used for GC/MS analysis.

GC/MS analysis was carried out using an AOC-6000 autosampler (Shimadzu Co., Kyoto, Japan) and a GCMS-8040 gas chromatograph/mass spectrometer (Shimadzu Co.) equipped with a BPX-5 capillary column (internal diameter: 30 m × 0.25 mm; film thickness: 0.25 µm; SEG, Victoria, Australia) as previously described [22]. Next, 20 µL of MSTFA was added to the sample supernatant, and the mixture was incubated at 750 rpm for 30 min at 37°C. The derivatized solution (1.0 µL) was injected into the GCMS-8040.

GC/MS analysis was conducted using the Smart Metabolites Database Ver. 2 (Shimadzu, Co.), which contains information regarding GC analytical conditions, MRM parameters, and retention index employed for the metabolite measurement. The raw data were processed using GCMSsolution and LabSolutions Insight, including noise filtering and peak processing.

Using the GC/MS method, 150 metabolites and 28 internal standards, including 27 isotope-labeled metabolites and 2-isopropylmalic acid, were analyzed. These metabolites and internal standards were grouped by hierarchical clustering based on their physical and chemical properties [23, 24]. One metabolite closest to the centroid within each cluster was selected, and its stable isotope-labeled compound was used as a surrogate. The peak area ratio was calculated by dividing the peak area of the metabolite by that of the metabolite surrogate in the same group and used for Workflow 1.

For quantitative analysis of the candidate biomarkers, a mixture of a metabolite surrogate of known concentration and one reference compound from each group was measured. Five mixtures were measured, in which the concentration of the metabolite surrogate remained constant while the concentration of the reference compound varied. Calibration curves were constructed from the peak area ratios. The lower limit of quantification (LLOQ) was determined based on information from public databases [25, 26], and five concentrations, at the LLOQ and at 10-, 50–, 75–, and 100–fold relative to the LLOQ, were established for the reference compound. For the other metabolites within the same group, the concentration was calculated from the calibration curve of the reference compound using their relative response factors with respect to the reference compound. The quantity of metabolite candidate biomarkers was confirmed similarly based on a calibration curve calculation for the reference compound. Eleven metabolites did not meet the ±50% accuracy criterion for concentration differences between the adjacent measurement points and were excluded from the 45 metabolite candidate biomarkers, which were selected by the Workflow 1 Data analysis. The detected metabolites, metabolite surrogates, and reference compounds are listed in S1 Table.

### Data analysis

#### Workflow 1: select features

Feature selection was conducted prior to model development. All performance metrics were estimated using independently split test datasets to reduce potential information leakage. The metabolite data were split into training and test data sets at a ratio of 7:3 by stratified random sampling after excluding exogenous compounds, unstable metabolites during storage and measurement, and highly correlated metabolites from pre-selection (Fig 2 (A)). Training dataset was scaled using autoscaling to re–scale each metabolite feature and then divided into five subsets. For each trial, the five subsets were used for 5-fold cross-validation to evaluate a set of randomly generated prediction parameters. The mean cross-validation score was used as the evaluation value for that trial. This process was repeated 50 times for internal validation, and the parameter set with the best average score was selected as the optimal set. Next, the estimated model was applied to the test data as external validation. This cycle was repeated 30 times. Because different ML algorithms may predict or rank different classification results, Lasso logistic regression (LR) [27], random forest classifier (RFC) [28], and support vector machine (SVM) [29] algorithms were used to select metabolite features for internal validation. Recursive feature elimination and variable importance in projection were used for feature selection for RFC and SVM, respectively. The importance of the metabolites in the classification model was determined by the coefficients of the model for LR, Gini impurity for RFC, and permutation feature importance for SVM in internal validation. The importance was adopted for the models in which AUC in the external validation was ≥ 0.7.

**Fig 2.**
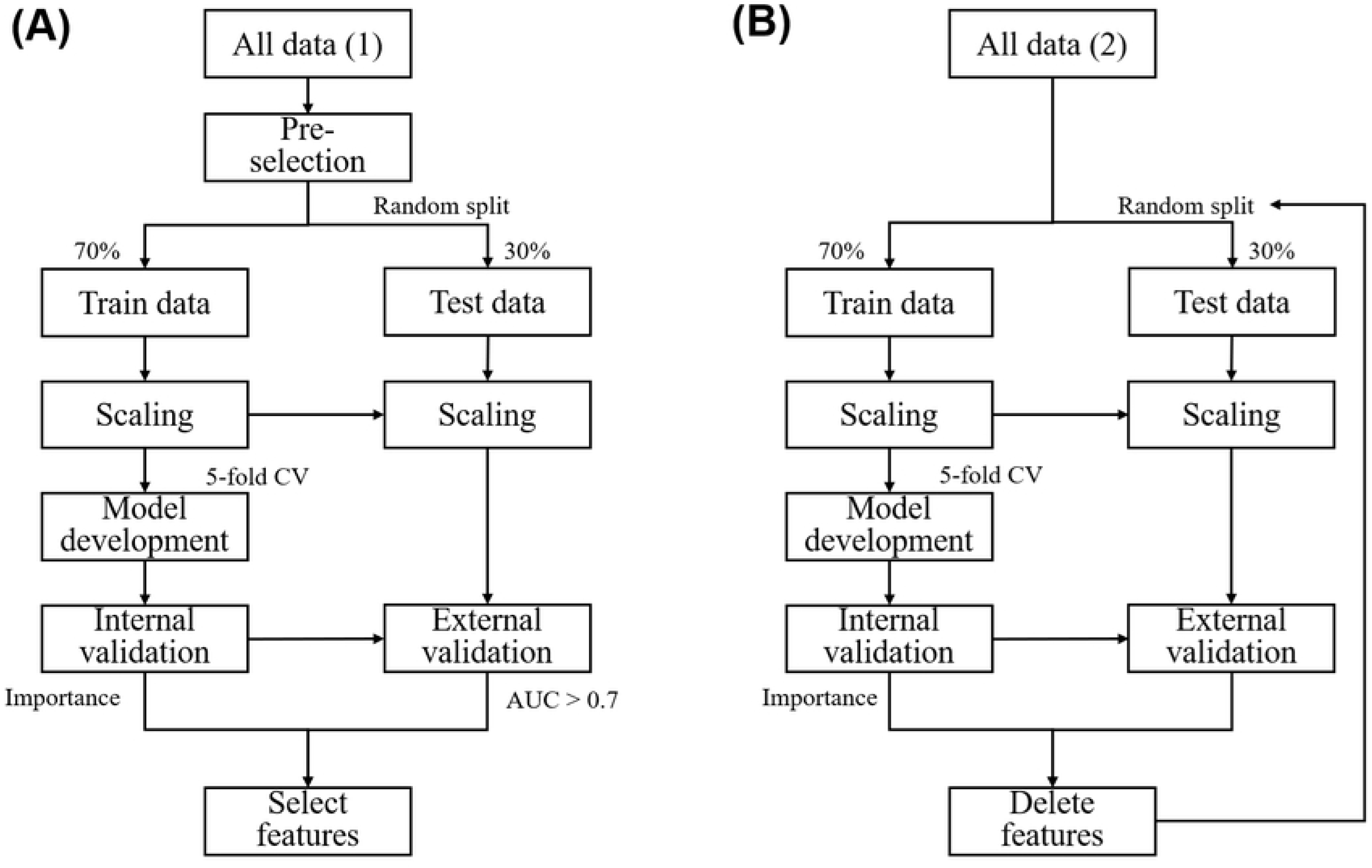
Data Analysis workflow. (A) Workflow 1: Select features. The data were randomly split into training and test sets. Machine-learning (ML) classification models were established using cross-validation (CV) of the training data set and validated using the test data set. The importance of each metabolite was calculated by the sum of the importance of the models, which revealed an AUC > 0.7 for the external validation with test data. (B) Workflow 2. Model development. The least important metabolite feature was sequentially removed, and the data were randomly split into training and test data sets at each iteration. The model was then redeveloped using the remaining metabolite features, and the classification performance was evaluated. This procedure was repeated across different numbers of metabolite features to identify the feature subset yielding the highest mean external validation AUC.

#### Workflow 2: model development

Model development was performed using independently split test data sets to reduce potential information leakage. As in Workflow 1, the data were randomly split into training and test data sets and used to develop classification models. To determine the optimal number of metabolites for each ML algorithm, a recursive feature elimination (RFE)-based strategy was applied [30]. Briefly, the least important metabolite feature was sequentially removed from those selected in Workflow 1, and the model was redeveloped at each iteration to assess its predictive performance across different numbers of metabolite features. The optimal number of metabolites was defined as the feature subset size that yielded the highest mean AUC in the repeated external validation [31].

#### Workflow 3: independent assessment using external HC samples

The models were fitted to the HC samples collected at another facility. The predicted probabilities between the subject groups and cancer stages were evaluated using Dunn’s all-pairs test for joint ranks. JMP Pro (ver. 18.2.0; SAS Institute Inc.) was used for the standard statistical tests, and Python (ver. 3.11.9; https://www.python.org/) was used for ML analysis.

## Results

### Select features

For the pre-selection step, we excluded 32 of 150 metabolites from the metabolite features based on four factors: exogenous compounds, unstable metabolites during storage of whole blood at 4°C, poor reproducibility, and highly correlated metabolites (S1 Table). The peak area ratio for 118 metabolites, which were selected during pre-selection, was calculated as described in “Plasma preparation and metabolomics analysis” section.

Fig 3 shows the importance of the metabolites in the classification models, which was calculated in the model development step of Workflow 1 and presented as a quartile plot. Filled circles represent the cumulative contribution ratio obtained by cumulatively summing the sum of importance ratios in descending order of the metabolites across 50 repeated CVs. The importance ratio was calculated by normalizing the importance by the sum of the importance of all metabolites. The metabolites are ordered from left to right by decreasing median importance, and those up to a cumulative contribution ratio of 80% are shown.

**Fig 3.**
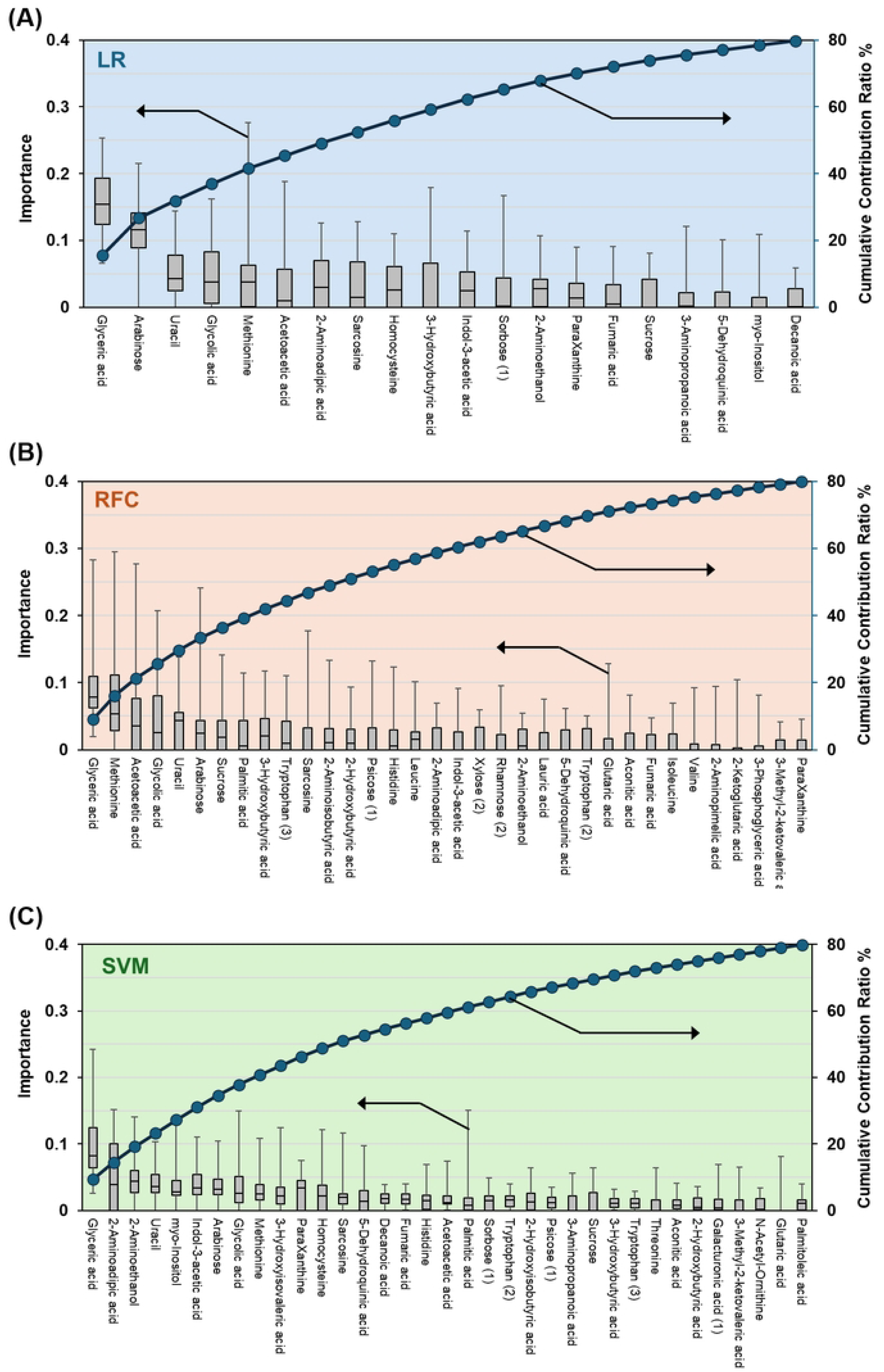
Importance of the metabolites estimated in Workflow 1 analysis with the LR (A), RFC (B) and SVM (C) algorithms. The importance of each metabolite is shown as a boxplot. Error bars indicate the minimum and maximum importances. Filled circles and solid lines indicate the cumulative contribution ratios of the metabolites. The left and right y-axes indicate the importance and the cumulative contribution ratio of each metabolite, respectively.

Glyceric acid, arabinose, uracil, glycolic acid, methionine, and acetoacetic acid exhibited higher importance and were consistently ranked among the top 10 across all ML algorithms. In contrast, 2-aminoethanol was ranked third in SVM, 13^th^ in LR, and 21^st^ in RFC, which indicates that metabolite importance in the classification model varied among the ML algorithms.

Metabolites contributing up to 80% of the cumulative contribution ratio in Fig 3 were selected as candidate biomarkers. A total of 45 metabolites, including 20 from LR, 34 from RFC, and 35 from SVM, were selected as plasma candidate biomarkers (S2 Table).

### Model development

The concentration of metabolite features was used in the Workflow 2 data analysis as described in “Plasma preparation and metabolomics analysis” section. The 34 metabolite quantitative data were used for model development. The metabolites used in three or fewer models were first excluded. The remaining metabolites were subjected to iterative feature elimination based on the summed importance ratios across the models. At each iteration, the least important metabolite was removed, and the model was redeveloped using the remaining features. This procedure was repeated until one feature remained for LR and two features remained for RFC and SVM. Fig 4 shows the number of metabolite features plotted against the mean AUC for the external validation across 30 repetitions. As the number of features was reduced, the mean external-validation AUC initially increased and subsequently decreased for all three algorithms. This indicates that model performance depended on an algorithm-specific balance between the information retained by the feature set and the risk of fitting sample-specific variation. The number of optimal features differed across algorithms: 15 for LR, 10 for RFC, and 16 for SVM, with corresponding mean AUCs in test data set of 0.877, 0.834, and 0.824, respectively. Because the optimal feature number differed across algorithms, and LR consistently achieved higher external validation AUCs compared with RFC and SVM at comparable feature numbers, suggest that performance differences cannot be attributed to feature dimensionality alone. Instead, they likely reflect differences in model capacity, regularization, decision boundaries, and the way each algorithm captured the signal and noise in the metabolomics data. The range of AUC values for the training and test data sets overlapped in the learning curves for three ML algorithms (S1 Fig), which indicates that overfitting was reduced by selecting the optimal metabolite feature set for each ML algorithm.

**Fig 4.**
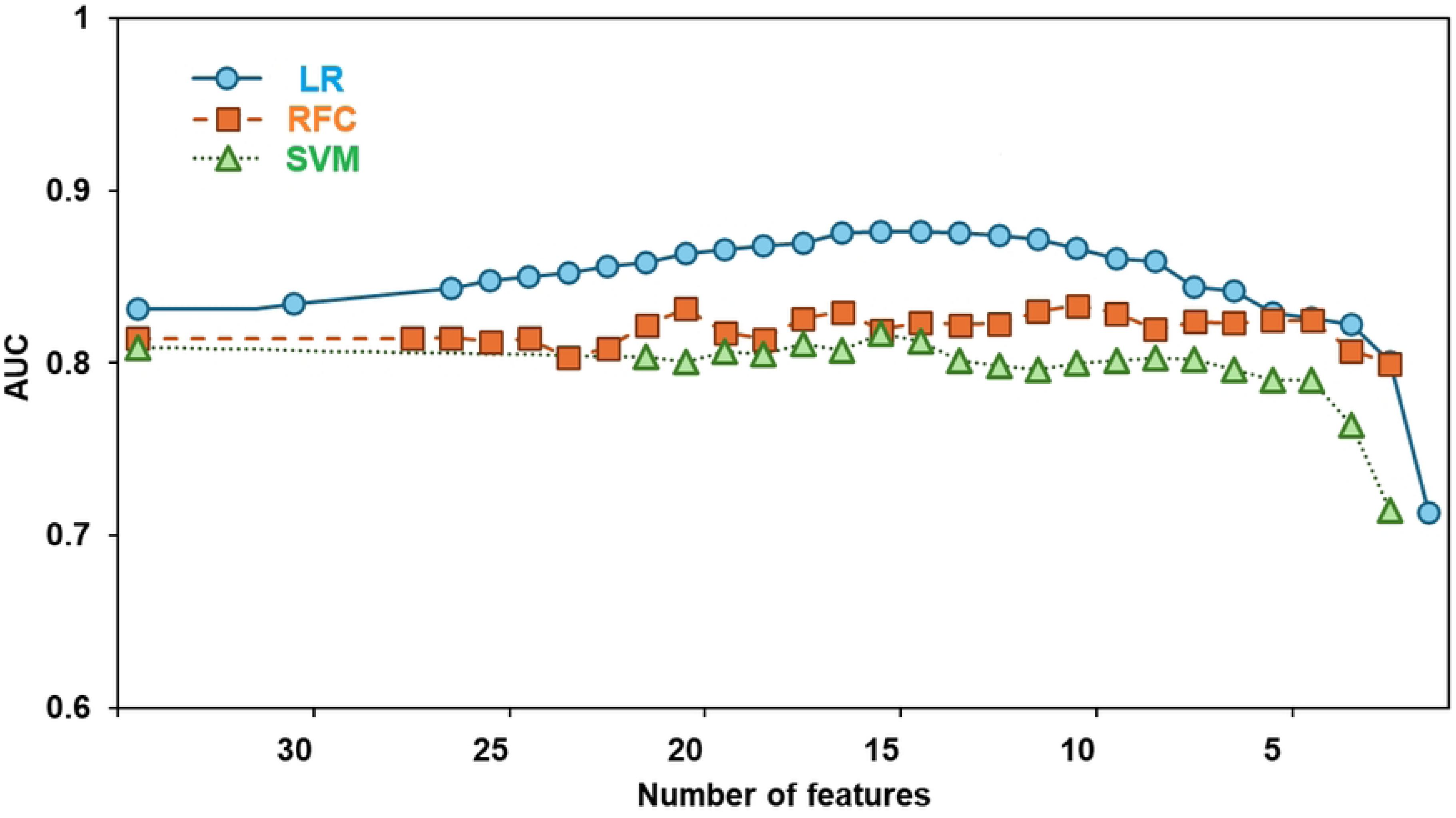
Relationship between AUC and the number of metabolite features in Workflow 2 model development. The average AUC of external CV validation for the classification models with three machine-learning algorisms LR (blue circles and bold line), RFC (orange squares and dashed line), and SVM (green triangles and dotted line), is shown.

### Assessment of classification models

The classification model with the smallest difference between the internal and external validation AUCs was selected as the best-performing model. Fig 5 shows the ROC curves for the training (A), test (B), and all (C) data sets for the best-performing models obtained by applying the metabolite features selected in “Model development” section to the three ML algorithms. The performance of the LR, RFC, and SVM models is summarized in S3 Table. In test data set of the three best-performing models, LR exhibited the highest AUC (0.910; 95% confidence interval (CI): 0.826–0.974), followed by RFC (0.893; 95% CI: 0.805–0.961) and SVM (0.843; 95% CI: 0.736–0.930). The differences in probability between BC and HC by the three MLs are shown in Fig 5 (D). HC2 and HC3 represent healthy controls collected from different facilities. Significant differences were observed between BC and HC2, whereas no significant difference between HC2 and HC3 was evident (Dunn’s post-tests of the Kruskal–Wallis test) in the models. The stage-specific differences in probabilities of BC are shown in S2 Fig. The values for stage III and IV were combined because of the small number of stage IV samples (n = 3). Significant differences were observed for all stages between HC2 in the models. The metabolite features used in the three best-performing models are listed in S4 Table. Importantly, several metabolites were consistently selected across different ML algorithms, suggesting that these candidate biomarkers reflect stable disease-associated signals rather than model-specific artifacts.

**Fig 5.**
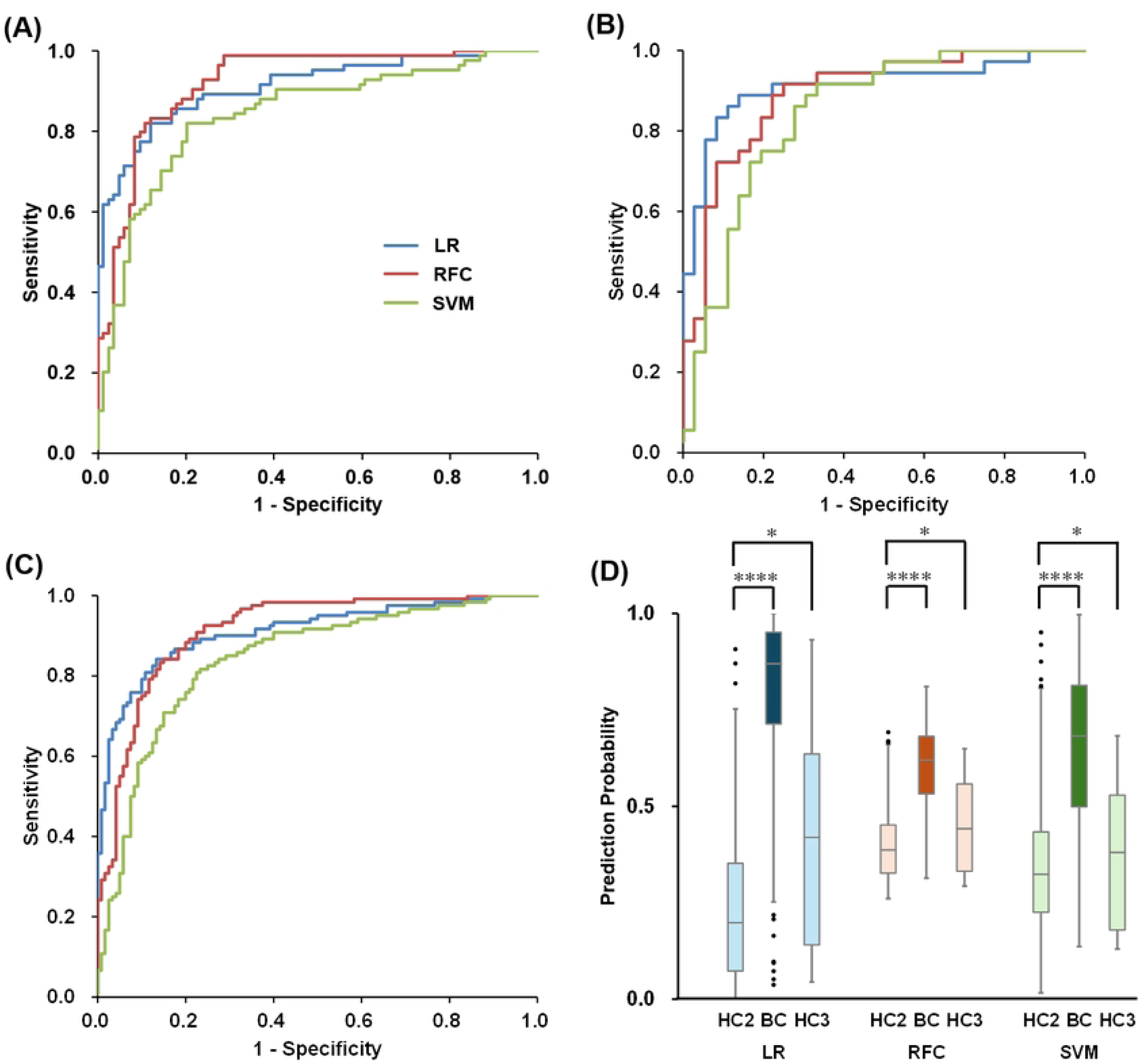
Discriminative performance of the classification models by three machine-learning (ML) algorithms. Receiver operating characteristic (ROC) curves for all data (A), with cross-validation (CV) results for the training and test data sets shown in (B) and (C), respectively, for LR, RFC, and SVM. Panel (D) shows the box plots for the prediction probabilities for HC2, breast cancer (BC), and HC3 using Workflow 2 and 3 data. Filled circles indicate the outliers in box plots. An outlier was defined as a data point that lies below Q1 − 1.5 × (Q3-Q1) or above Q3 + 1.5 × (Q3-Q1), in which Q1 and Q3 are the first and third quantiles. In (D), asterisks indicate the p-value of Dunn’s post-test after the Kruskal–Wallis test. * p > 0.05 and **** p < 0.0001.

## Discussion

The principal contribution in the present study is the establishment and evaluation of a plasma metabolomics workflow along with the introduction of conventional ML algorithms. The workflow integrates controlled prospective sample collection, quantitative GC/MS measurement, analytical quality-based feature filtering, selecting ML algorithm-specific optimal metabolite feature sets using a recursive feature elimination-based strategy, repeated external validation using a split test data set, and assessment in healthy controls collected at another facility. This design addresses several translational barriers that have limited the clinical implementation of plasma metabolomics, including preanalytical variability, analytical reproducibility, overfitting, and the lack of independent assessment. The use of Lasso logistic regression, random forest classifier, and support vector machine was intentional. These are established algorithms that allow the contribution of the data quality and validation framework to be evaluated without attributing performance to an untested or highly specialized modeling method. Comparing the three best-performing classification models, the regularized LR model yielded the highest AUC (0.910; 95% CI: 0.826–0.974), followed by the RFC model (AUC = 0.893; 95% CI: 0.805–0.961) and SVM model (AUC = 0.843; 95% CI: 0.736–0.930). The optimal feature numbers differed across algorithms, and LR consistently outperformed RFC and SVM at comparable feature numbers. These results indicate that the observed performance differences are not attributable to feature dimensionality alone, but also reflect how each algorithm represents the metabolomic signal and controlled model flexibility. To confirm whether the high AUCs of the classification models were primarily contributed by the intrinsic structure of the original metabolic data, optimization of the number of features, or the model-learning process itself, we performed principal component analysis (PCA) as an unsupervised analysis. Next, the resulting principal component scores were used for k-nearest neighbors (k-NN) classification, and the AUC, accuracy, sensitivity, and specificity were evaluated (S3 Fig and S5 Table). PCA based on the 34 metabolites selected in Workflow 2 did not separate HC from BC, and all data were uniformly distributed without bias or distinct clustering. When k-NN analysis was conducted using the PCA scores, the AUC was low (0.593), suggesting that differences in the intrinsic structure of the original data contributed little to the high AUC values obtained by the three best-performing models. Similarly, PCA was conducted using the features selected by LR, RFC, and SVM. None of the three PCA analyses revealed a clear separation between the HC and BC data, and the k-NN analyses did not show a substantial improvement in classification performance. These results suggest that the high AUCs were not likely to be explained solely by changes in the data structure resulting from the optimization of the features. Taken together, the high AUCs achieved by the three best-performing models may have resulted from the combination of subtle differences in the data structure and the intrinsic characteristics of the ML algorithms. The structure of the plasma metabolomics data may be expressed relatively linearly and additively, because LR is a linear classification method that directly estimates coefficients, whereas Lasso regularization reduces unnecessary coefficients [27]. To leverage the complementary advantages of each algorithm, we constructed a bootstrap aggregating (bagging) ensemble model. The bagging model achieved a balanced and superior performance with a high AUC (0.911; 95% CI: 0.872–0.946), as shown in S4 Fig, stable sensitivity (68.3%), and a low Brier score (0.136), while yielding only a single false positive in the independent healthy control (HC3) dataset. This suggests that an ensemble approach will be useful for future applications, such as a clinical triage or screening test.

The importance of the three best-performing models provides information beyond the selection of an optimal feature number. Because the algorithms ranked metabolites differently, features selected by only one algorithm may partly reflect model-specific representations of the data. In contrast, metabolites that were consistently selected or assigned high importance across LR, RFC, and SVM are less dependent on the particular modeling choice and may represent more robust disease-associated signals. The fold-change (FC) of the metabolites selected by the ML algorithms and their importance within models are shown in Fig 6 and S4 Table. The metabolite candidate biomarkers were ranked in descending order based on the sum of their feature importances across the three best-performing models. The importance of each metabolite candidate biomarker was not directly correlated with its FC value. These results suggest that selecting a metabolite feature dataset with a data structure suited to the characteristics of each ML algorithm is more important for developing stable and high-performance binary classification models, rather than selecting biomarkers solely on the basis of significant differences between two classes [32].

**Fig 6.**
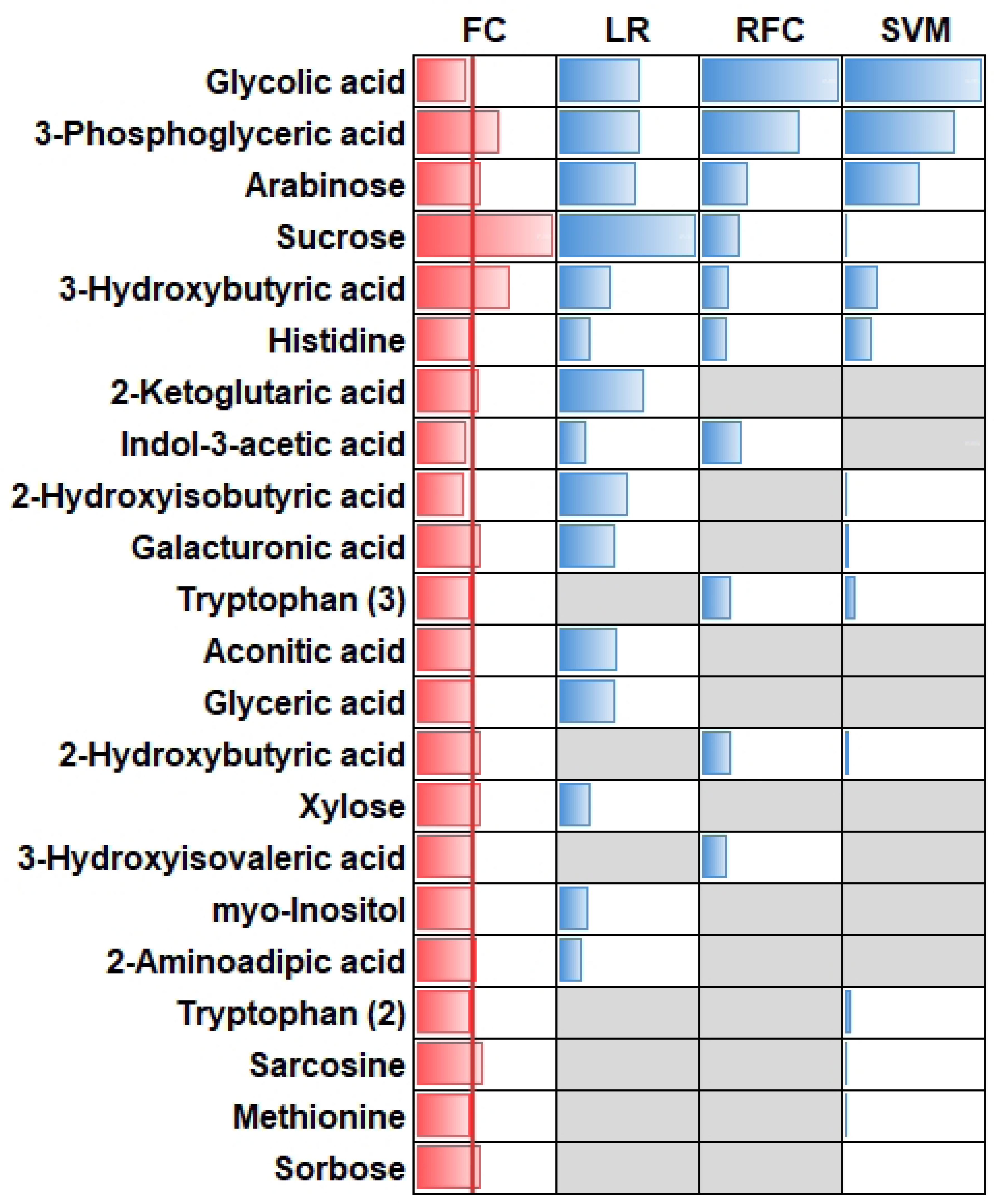
Fold change (FC) and feature importance of each metabolite feature in the best-performing models by three ML algorithms. The red bars indicate FC, and the blue bars indicate the importance of the metabolites. The solid red line in the FC column indicates FC =1. Grey cells indicate that the corresponding metabolite was not used in the classification model.

Furthermore, compared with distinguishing HC from BC using a single metabolite [33], constructing a classification model with an ML algorithm based on data from multiple metabolites, including those that do not show statistically significant differences between the two groups [34], may result in a more robust and stable model. In our best-performing models, five core metabolites, 3-hydroxybutyric acid, 3-phosphoglyceric acid, glycolic acid, arabinose, and histidine, exhibited persistently high variable importance across all three ML algorithms, indicating they captured stable disease-associated signals independent of the model architecture. Based on previous studies [13, 34–36], these core candidate biomarkers consistently selected across the multi-algorithm pipeline suggest coordinated systemic alterations in host–tumor interactions and energy homeostasis. These results may lead to novel treatments for BC in the future. Thus, algorithmic concordance provides a complementary criterion for prioritizing candidate biomarkers, although it does not replace validation in independent BC and benign-disease cohorts.

This study has several limitations that should be acknowledged before considering clinical implementation. First, the independent external dataset included exclusively healthy controls (HC3) and lacked an independent breast cancer cohort. Consequently, while the workflow demonstrated high stability in reducing false positives, the true diagnostic sensitivity of the optimized models remains unverified in an external patient population. Second, potential confounding variables, including patient body mass index, systemic metabolic comorbidities, menopausal status, and current medications, were not controlled or stratified. Third, the case– control study design inherently carries a risk of spectrum bias, which may overestimate diagnostic performance relative to real-world screening populations, in which the prior probability of disease is substantially lower. Finally, no direct comparisons or cross-validations were conducted against benign breast diseases (e.g., fibroadenomas) or other malignancy types, leaving the disease-specificity of this metabolic signature yet to be established.

## Conclusions

Our multi-algorithm standardized workflow successfully identified a robust plasma metabolomic signature capable of stable discrimination between BC and HC within this study population. However, to establish its broader clinical applicability, multi-center prospective validation incorporating independent cancer cohorts and benign disease controls is fundamentally required.

## Acknowledgments

The authors thank the members of the Department of Breast and Endocrine Surgery, Health and Medicine Clinic, and Umeda Health and Medicine Clinic of Hyogo Medical University for collecting plasma samples. The authors thank Reiko Tanaka, Konomi Fukui, Maya Yamazaki, Miku Shimizu, Rina Fujita, Chiyomi Hamada, and Keiko Wakai for measuring plasma samples, and thank Shota Oshikawa and Shigeki Kajihara for supporting data curation.

## Supporting information

**S1 Fig. Learning curves for the LR (A), RFC (B), and SVM (C) models.** Blue and red circles and shaded bands indicate the learning curves for cross-validation of the training and test data sets, respectively. Circles represent average AUC values, and the upper and lower boundaries of the bands indicate the maximum and minimum AUC values, respectively.

**S2 Fig. Prediction probabilities for HC2, each stage of breast cancer (BC), and HC3 using Workflow 2 and 3 data.** Panels A, B, and C show the results of LR, RFC, and SVM, respectively. Filled circles and asterisks in B indicate the outliers and p-values, which are the same as in Fig 5 (D).

**S3 Fig. Results of Principle component analysis (PCA).** PCA score plots with all 34 features (A). Blue and red circles show HC2 and BC, respectively. The blue and red lines indicate the 95% confidence ellipses for HC and BC, respectively. The PCA score plots with the features that were used in the best-performing LR, RFC, and SVM ML models are shown in (B), (C), and (D), respectively.

**S4 Fig. Discriminative performance of the machine-learning (ML) bagging model.** Receiver operating characteristic (ROC) curves for all data (A). AUC values are presented with a 95% confidence interval (CI) in parentheses. The prediction probability (B) for HC2, breast cancer (BC), and HC3 using Workflow 2 and 3 datasets. Filled circles and asterisks in B indicate the outliers and p-values, the same as in Fig. 5 (D).

**S1 Table. Grouping of Metabolites and Internal standards in Workflow 1 and 2.**

**S2 Table.** The **Cumulative contribution ratio % of selected metabolites in Workflow 1.**

**S3 Table. Performance of binary classification models by three machine-learning algorithms.**

**S4 Table. Fold-change, BC/HC, and the importance of metabolite candidate biomarkers in the best-performing classification model based on three ML algorithms.**

**S5 Table. Results of PCA and k-NN analysis with selected metabolite features.**

